# MMP2 loss leads to defective parturition and severe dystocia in mice

**DOI:** 10.1101/2022.11.22.517326

**Authors:** Rotem Kalev-Altman, Tamar Levy, Nahum Y. Shpigel, Efrat Monsonego-Ornan, Dalit Sela-Donenfeld

## Abstract

Parturition is the final step of mammalian reproduction and an essential process for the species’ survival. During pregnancy, the uterus is maintained quiescence which is important for fetal growth and development. However, at term, fundamental changes in myometrial contractility are initiated for efficient expulsion of the fetus. These changes involve tissue remodeling that requires changes in the extracellular matrix (ECM). The gelatinases subgroup of matrix metalloproteinases (MMPs), has only two members: MMP2 and MMP9, which are both known to participate in uterine ECM remodeling throughout the estrus cycle as well as during pregnancy, parturition and postpartum involution. Yet, no knowledge exists regarding their loss-of-function impact on the uterus. Here we investigated the effect of MMP2 and/or MMP9 genetic loss on parturition process. Single and double knockout (dKO) mice for MMP2 and/or MMP9 were used. We found high percentages of dystocia in mmp2^-/-^, mmp2^-/-^mmp9^+/-^ and dKO females, but not in mmp9^-/-^ females. Histological analysis of nulliparous uterine tissue of WT, mmp2^-/-^, mmp9^-/-^ and dKO, at 8 weeks, 4 months and 8-9.5 months, revealed that the uterine tissue of mmp2^-/-^ presents alterations in tissue size and structure, mainly when reaching to 8-9.5 months of age, including enlarged total tissue, myometrium, endometrium and luminal cavity. Additionally, Masson’s Trichrome staining suggested a mechanism of extensive fibrosis in mmp2^-/-^ myometrium, which may lead to dystocia. Altogether, our research highlights a novel cause for dystocia pathology mediated by loss of MMP2 activity in uterine tissue during mammalian parturition.

## Introduction

Parturition is the final step of mammalian reproduction, an essential process for the successful production of the next generation. During pregnancy, the uterus undergoes a period of quiescence which is important for fetal growth and development. Yet, at some point, dramatic changes occur in myometrial contractility that result in the efficient expulsion of the fetus. These events are necessary for the survival of both the fetus and the mother. However, the nature of these pathways is only partially defined and increased knowledge of the cascade of events that occur at parturition will lead to advances in combating preterm births or optimization of protocols for medically induced labor [1]. Preterm births count for ∼11.5% of all live births in the US and are the main cause of perinatal mortality and morbidity worldwide [2,3]. Moreover, based on the world health organization (WHO), up to 25% of all deliveries at term in developed countries, involve the artificial induction of labor. However, it is unclear whether the need to induce the labor is due to difficulties in parturition or other reasons.

The uterine wall consists of three major elements: (1) the endometrium which contains multiple secreting glands that undergo extensive remodeling throughout the estrus cycle [4]; (2) the myometrium which consists of an inner circular layer and an outer longitudinal layer of oriented smooth muscle, and (3) the perimetrium outer layer. In mice, the estrus cycle is short (4-5 days) and compromises four precise phases including proestrus, estrus, metestrus, and diestrus. When mice are being used as an experimental model in reproductive biology, it is crucial to assess the functioning status of the female reproductive system by estimating or synchronizing the estrus cycle [5–7].

The uterine transition from quiescence to contraction is a mandatory shift for executing natural parturition. Research from past decades has provided critical insights into the parturition process, but the molecular mechanisms involved in its initiation remain largely unknown. The myometrium layer of the uterine wall plays an essential role in regulating uterine quiescence and contraction by secreting and/or responding to progesterone and estrogen signaling, which in turn regulate the myometrium activities [2,3]. Furthermore, similar to other muscle cell contractions, myometrial contractions are also mediated by elevated intracellular calcium concentration (Ca^+2^), which is regulated by both the release of Ca^+2^ from intracellular stores in the sarcoplasmic reticulum and Ca^+2^ entry from the extracellular space. It is well-studied that both sarcoplasmic reticulum Ca^+2^ efflux and extracellular Ca^+2^ influx in myometrial smooth muscle cells, are important for myometrial contractions during pregnancy [8,9]. However, the molecular mechanisms regulating Ca^+2^ mobility in the myometrium and uterine contractility during parturition are not fully understood.

Matrix metalloproteinases (MMPs) are a large family of enzymes known for their ability to degrade different components and proteins in the extracellular matrix (ECM). As such, they were found to be involved in many physiological processes, such as in ovulation and embryo implantation; but also in pathological conditions, including cancer cell metastasis, Multiple Sclerosis and Alzheimer’s disease [10–14]. The gelatinases subfamily of MMPs, has only two members, MMP2 and MMP9. These two secreted enzymes were described previously in the literature to have a similar structure and active site, a comparable biological activity in different biological systems, and the same substrates *in vitro* [15–17]. Concomitantly, we have previously found that during development, both MMP2 and MMP9 are required for executing the migration of the unique population of neural crest cells (NCCs) in both chick and mouse embryos, and each MMP compensates for the loss of the other [18]; see also Duong and Erickson 2004 and Monsonego-Ornan et al., 2012 [19,20].On the other hand, in our recent study on skeleton development, we found that each gelatinase has an individual and unique role; while MMP2 was found to have a role in intramembranous ossification affecting the development of the skull and the cortices, MMP9 was demonstrated to have a role in endochondral ossification affecting the longitudinal growth of the skeleton, and the overall body length [21]. Hence, their redundant or individual activities are highly context dependent.

Interestingly, several MMPs, including MMP1, MMP2, MMP3, MMP7, MMP9, MMP10 and MMP11, and their inhibitors, the tissue inhibitors of MMPs (TIMPs), including TIMP1, TIMP2 and TIMP3, were previously found to be expressed in the uterus of various menstruating and non-menstruating species [22]. Furthermore, their balanced function was shown to be essential for normal uterine tissue remodeling throughout the estrus cycle as well as during pregnancy, parturition and postpartum uterine involution in menstruating and non-menstruating species, such as rodents and humans [22–24]. These studies suggested that MMP family members participate in the regulation of the uterus dynamic remodeling. Yet, the comparable role of each gelatinase sub-member has not been explored. In the present study, we set to specifically determine whether the gelatinases, MMP2 and MMP9, have a combined or individual role in uterine function during parturition process in mice. By studying single and double-KO (dKO) mice for MMP2 and/or MMP9, we found extremely high percentage of defective parturition process which led to high rates of dystocia in mmp2^-/-^, but not mmp9^-/-^, pregnant females. Histological analysis of uteri from nulliparous females of different MMP2/MMP9 null genotypes demonstrated distorted uteri in MMP2-null females. To the best of our knowledge, this is the first description of the effect of single or double loss of MMP2 and/or MMP9 genes on uterine function during physiological parturition process in mice.

## Material and methods

### Animals

Mmp2^-/-^ [25,26], mmp9^-/-^ [27], dKO (mmp2^-/-^mmp9^-/-^) [21] and wild-type (WT) C57BL/6J mice were used. WT and mmp9^-/-^ mice were purchased from Harlan laboratories (Rehovot, Israel), while mmp2^-/-^ mice were provided from Martignetti Lab (Mount Sinai School of Medicine, NY, USA). dKO mice were generated at our lab, as detailed in Kalev-Altman 2020, and Kalev-Altman 2022 [18,21]. Additional genotypes (mmp2^+/-^, mmp9^+/-^, mmp2^+/-^mmp9^+/-^, mmp2^-/-^mmp9^+/-^, mmp2^+/-^mmp9^-/-^) were generated at the lab. Mice were maintained at the Hebrew University Specific Pathogen Free animal facility according to animal care regulations. All procedures were approved by the Hebrew University Animal Care Committee (license number 21-16781-3).

### Genotyping

Genotyping was performed as was previously described [18]. Briefly, DNA was extracted from an ear tissue, genotyped by PCR (Biometra, Germany) and analyzed in 2% agarose gel. For each mutated gene (*mmp2* or *mmp9*), primers were designed by choosing one primer inside the neomycin insert and another downstream to the neomycin insert in order to specifically recognize each MMP-mutated gene [18].

### Genotype distribution calculation and litter size measurements

For calculating the genotypic distribution, males and females of mmp2^+/-^ genotype were mated and females were monitored for vaginal plug which was determined as gestational day (GD) 0.5. 18 days later, at GD18.5, pregnant females were sacrificed and their fetuses were harvested. A small tissue from each fetus was taken for genotyping. The relative distribution of each possible genotype (WT, mmp2^+/-^, and mmp2^-/-^) was calculated and compared to the expected ratio according to Mendel’s law for genetic distribution.

For litter size measurements, 37 pairs of WT mice and 11 pairs of mmp2^-/-^, mmp9^-/-^ and dKO mice were mated and the number of neonates was counted at birth (postnatal day 0, P0).

### Collection of uterine tissue from synchronized females

Nulliparous females were euthanized at the age of 8 weeks, 4 months and 8-9.5 months and their uterine tissue was collected. For the collection of mouse tissues under synchronization of their hormonal cycle, mice were treated with Depo-Provera (Pfizer Inc., New York City, NY, USA) which contains medroxyprogesterone acetate (MPA), for synchronization into diestrus stages. Mice were subcutaneously injected twice, on day 1 (starting day) and day 5, at the dose of 3 mg MPA per mouse. Synchronized diestrus stage was reached 7 days after first Depo-Provera administration [28]. When indicated, uteri at 12M of age were collected without synchronizing the estrus stage. Uterine morphology were grossly assessed, followed by preparation for histology.

### Histology

Uterine tissue was fixed with 4% paraformaldehyde (PFA) for overnight at 4°C, washed three times in PBS, dehydrated with increased concentrations of ethanol (70%, 80%, 90%, 2×100%), and transferred twice into xylene for 1 hour each. Samples were then incubated three times in liquid paraffin for 1 hour each at 58°C, and left to solidify at RT. Transverse tissue sections of 5µm were prepared with Leica microtome (Leica, Germany). For staining, slides were washed twice for 10 minutes in xylene for paraffin removal and rehydrated by 10 minutes washes in decreasing ethanol concentrations (2×100%, 90%, 70%, and 50%) and PBS.

H&E staining was conducted as described previously [29–31]. Masson’s trichrome staining was conducted as described before [32]. Briefly, slides were stained in Weigert’s Iron Hematoxylin solution, followed by rinsing in running warm tap water, and distilled water (dw). Next, slides were stained in Biebrich Scarlet-Acid Fuchsin solution and washed in dw. Then incubated in phosphomolybdic-phosphotungstic acid solution and transferred directly to aniline blue solution, followed by a quick rinse in dw and 1% acetic acid solution.

### Imaging

Stained sections were viewed by the light microscopy eclipse E400 Nikon, using light filters or by Axio Imager M1 microscope with AxioCam MRm camera (Zeiss, Germany), or scanned using Panoramic Flash III 250 scanner (Dhistech3).

### Quantification and statistical analysis

All data are expressed as the mean ± SD (standard deviation). The significance of differences between groups was determined using JMP 15.0 Statistical Discovery Software (SAS Institute 2000) by one-way ANOVA followed by the Tukey–Kramer HSD test and t-test. Differences were considered significant at P ≤ 0.05. Groups with the same letter were not measurably different, and groups that were measurably different are indicated by different letters. Groups may have more than one letter to reflect the “overlap” between the sets of groups, and sometimes a set of groups is associated with only a single treatment level.

## Results

### *Mmp2*-loss results in defective parturition process and dystocia

Dystocia is defined as a difficult labor and delivery with bad outcome. When breeding the different MMP2/MMP9-null colonies, we occasionally observed pregnant females with such phenomenon. Hence, we decided to methodologically characterize the occurrence of dystocia in the following genotypes: WT, mmp2^+/-^, mmp2^-/-^, mmp9^+/-^, mmp9^-/-^, mmp2^+/-^ mmp9^+/-^, mmp2^-/-^mmp9^+/-^, mmp2^+/-^mmp9^-/-^ and mmp2^-/-^mmp9^-/-^ (dKO); comparing to WT. Dystocia was defined as a condition in which a pregnancy has reached to 22.5 gestational day (GD) according to plug observation, and did not result in a normal parturition. In these cases, all females still had one or more (dead) fetuses in their uterus. Other cases were also considered as dystocia, such as vaginal bleeding at 18.5-21.5 GD or cases where GD was not determined by vaginal plug but it was clear by eye that the female had reached to the end of the pregnancy but parturition did not start, with no newborns at the cage and remaining of dead fetuses in-uterus (as observed by eye) (Fig. 1 A-C). In all of these cases the females were immediately euthanized.

**Figure 1.**
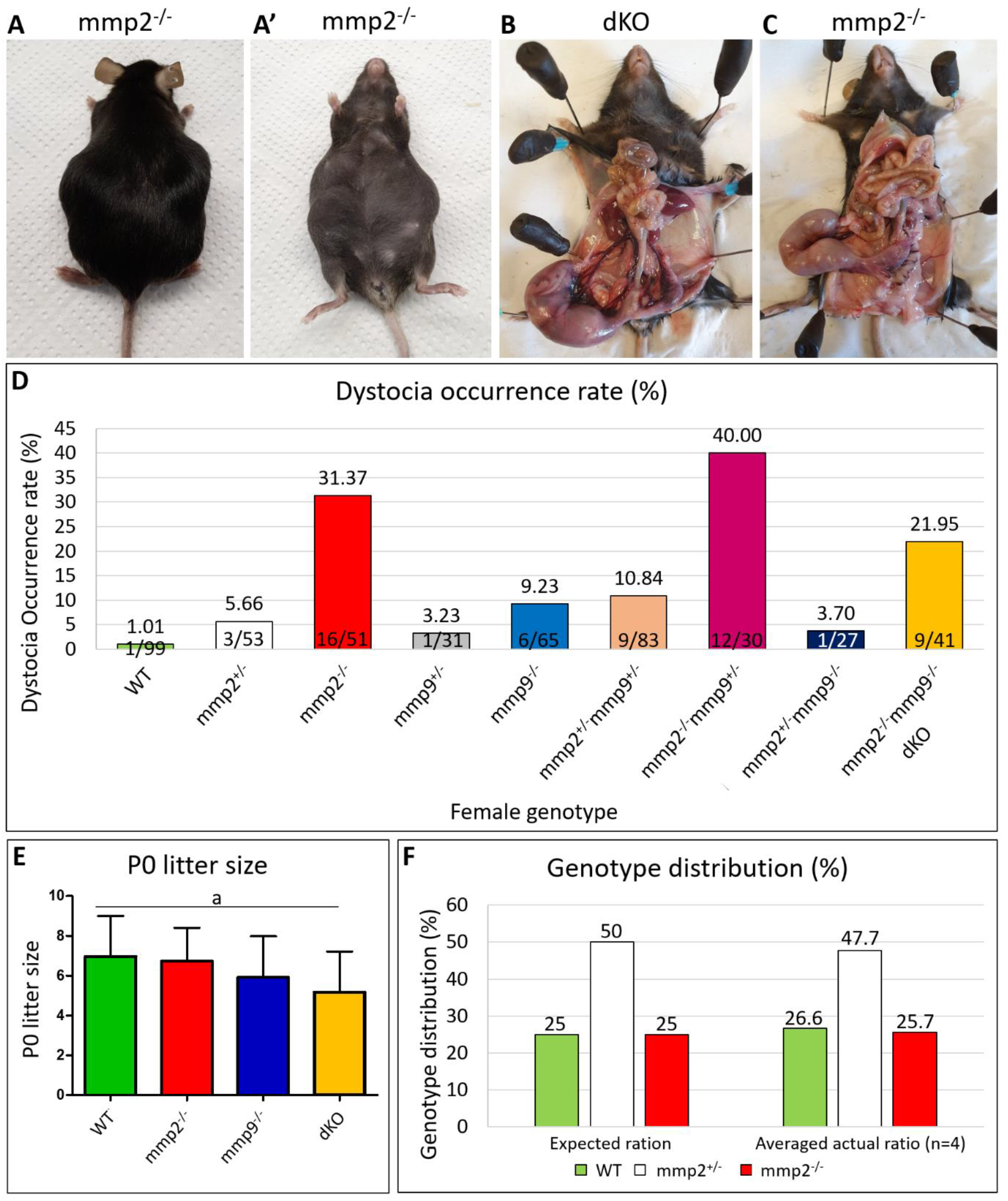
High percentage of dystocia in mmp2^-/-^, mmp2^-/-^mmp9^+/-^ and dKO females. (A) Representative example of mmp2^-/-^ female which underwent dystocia at the end of the pregnancy. (B,C) When abdomen is opened, one or more fetuses were found dead in the uterus. (D) Dystocia percentage according to MMP2 and/or MMP9-KO genotypes demonstrating its higher occurrence in mmp2^-/-^, mmp2^-/-^mmp9^+/-^ and dKO females (red, claret and orange columns, respectively), compared to the other genotypes. (E) Litter size measurements at P0 show no difference in mmp2 and/or mmp9 KO litter size compared to WT; n=37 for WT or n=11 for each KO. (F) Genotype distribution from mating mmp2^+/-^ female and male show no difference in generation of mmp2^-/-^ embryos (n=4 litters).

Dystocia percentages were measured for each genotype as the number of pregnancies ended in dystocia in relation to the total number of pregnancies for the same genotype measured over a period of 7 years (2015-2021). We found that a high percentage of 31.37% of all mmp2^-/-^ pregnancies (16/51) resulted in dystocia (Fig. 1 D). This high ratio is in great contrast to only 1.01% (1/99) or 9.23% (6/65) pregnancies that resulted in dystocia in WT and mmp9^-/-^ genotypes, respectively (Fig. 1 D). Moreover, measuring the dystocia percentage among females of double mutation in both MMP2 and MMP9 genes, such as double heterozygote mmp2^+/-^mmp9^+/-^, mmp2^-/-^mmp9^+/-^, mmp2^+/-^mmp9^-/-^ or double full knockout (mmp2^-/-^mmp9^-/-^, dKO), revealed varied results, such as 10.84% (9/83), 40% (12/30), 3.7% (1/27) and 21.95% (9/41), respectively (Fig. 1 D). Intriguingly, this data shows that while the additional loss of one allele of *mmp9* to the MMP2-KO (mmp2^-/-^mmp9^+/-^) leads to a significant increase in dystocia incidence (40%), compared to dystocia incident found in the single MMP2-KO (mmp2^-/-^) which occurred in 31.37% of pregnancies, the loss of both *mmp9* alleles on the background of MMP2-KO (dKO) leads to a reversed phenotype of a decrease in dystocia incidence (21.95%), relatively to the mmp2^-/-^ or mmp2^-/-^mmp9^+/-^ females (Fig. 1 D). Together, this analysis demonstrates that the occurrence of dystocia is largely linked to a loss of MMP2. As well as indicates that the additional effect of the loss of the MMP9 gene is dosage-depended, which either worsen or reduces the dystocia rate as found in the single MMP2-KO.

Next, we went to evaluate whether the affected females could not undergo normal parturition process due to in utero fetal death. Viability of fetuses and litter size were measured at GD18.5 and P0, respectively. Regardless of the genotype, the entire litter was found to be alive at GD18.5 (data not shown) and with a similar number of fetuses at P0 (6.97-5.18), although a tendency to a slightly smaller litter size was found in the dKO, but this was not statistically significant (P=0.0508, Fig. 1 E). Furthermore, when we measured the genotype distribution of WT, mmp2^+/-^, and mmp2^-/-^ in the neonates generated from the mating of mmp2^+/-^ females and males, we found that the averaged ratio of all litters examined was as expected by the Mendelian inheritance: 26.6%, 47.7% and 25.7% for WT, mmp2^+/-^ and mmp2^-/-^, respectively (Fig. 1 F). Together, these results rule out any association between fetal death and dystocia in the various MMP2/MMP9-null genotypes.

### Mmp2^-/-^ nulliparous females present relatively shorter uterine horns

After unraveling the connection between *Mmp2* genetic loss and dystocia occurrence in mice, we next set to determine the effect of *Mmp2*-loss on the morphology of the uterus. For that, we collected synchronized uteri from 4-12 nulliparous females from the following genotypes: WT, mmp2^-/-^, mmp9^-/-^ and dKO; at three different ages: 8 weeks (w), 4 months (M) and 8-9.5M. The decision to investigate nulliparous mice at different ages was due to the observation that dystocia incidents occur mostly among older females; out of 56 females that underwent dystocia, only 3 were aged less than 4M and up to 44 were aged more than 6M (data not shown). Moreover, the synchronization of the females was important for preventing the occurrence of natural morphological changes that are a result of the estrus cycle rather than of the studied genotypes. After harvest, the length of the uterus horn was measured and was found to be somewhat shorter in mmp2^-/-^ samples compared to WT, at all examined ages (Fig. 2); however, this difference was not statistically significant (P=0.0935, P=0.1991 and P=0.33 for WT *vs* mmp2^-/-^ at 8w, 4M and 8-9.5M, respectively). Additionally, H&E staining of sections taken from the different uteri demonstrated that some mmp2^-/-^, mmp2^-/-^mmp9^+/-^ and dKO samples taken at 12M of age were found to be markedly abnormal with extensive luminal area and secretary glands, compared to WT (Fig. 2, P). These tissues also demonstrated very thin myometrial layer and almost no endometrium stroma compared to WT’s (Fig. 2, P).

**Figure 2.**
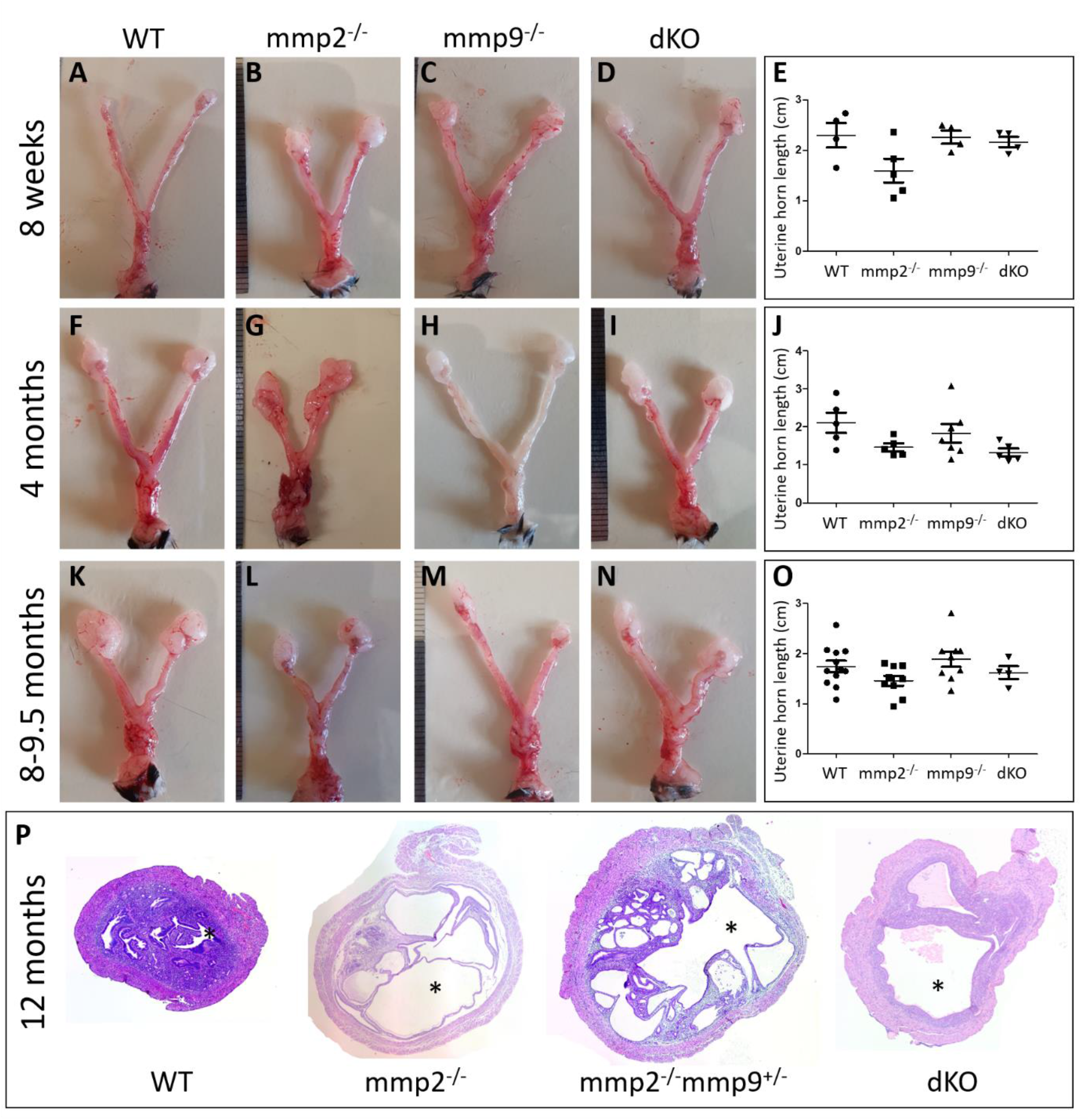
mmp2^-/-^ uteri are shorter in size, compared to WT. Representative photos of 8 weeks old (A-D), 4 months old (F-I) and 8-9.5 months old (K-N) diestrus stage uteri collected from all 4 genotypes: WT, mmp2^-/-^, mmp9^-/-^ and dKO. Measurements of the uterine horn length demonstrated that mmp2^-/-^ were relatively shorter at all ages (E, J, O), but not significantly. (P) Nulliparous uterine cross-section of mmp2^-/-^, mmp2^-/-^mmp9^+/-^ and dKO, show significant enlargement of the lumenal area and very thin myometrial layer (compare the lumen indicated by the asterisks).

### Mmp2^-/-^ uterus demonstrates enlargement of the myometrium, endometrium and lumen

To further determine whether the pathophysiology of the dystocia is associated with abnormal microscopic anatomy of the uterus at older stages, we conducted histological analysis on transverse sections from nulliparous uterine tissue at the age of 8w, 4M and 8-9.5M (Fig. 3), similar to the ages analyzed at Fig.2 A-O. The data revealed that at 8w and 4M, the uterine tissue of mmp2^-/-^, mmp9^-/-^ and dKO uterine presents a normal appearance compared to WT, as determined by the overall tissue size and the relative area taken by each layer of the uterine wall (Fig. 3 A-H). Yet, at 4M, the mmp2^-/-^ tissue starts to display a dissimilar structure with increased number of secreting glands in the endometrial sub-tissue, compared to other genotypes (Fig. 3 E-H). However, at the age of 8-9.5M, a significant enlargement of the entire tissue area was found in mmp2^-/-^ compared to other genotypes (Fig. 3 I-L). Furthermore, specific area measurements of each tissue type, such as the myometrium, endometrium, and the lumen, demonstrated that while at 8w and 4M no differences were found between the different genotypic groups, at 8-9.5M the area of all three measured elements (myometrium, endometrium and lumen) were significantly enlarged in mmp2^-/-^ compared to WT (Fig. 3, U-X compared to M-P and Q-T).

**Figure 3.**
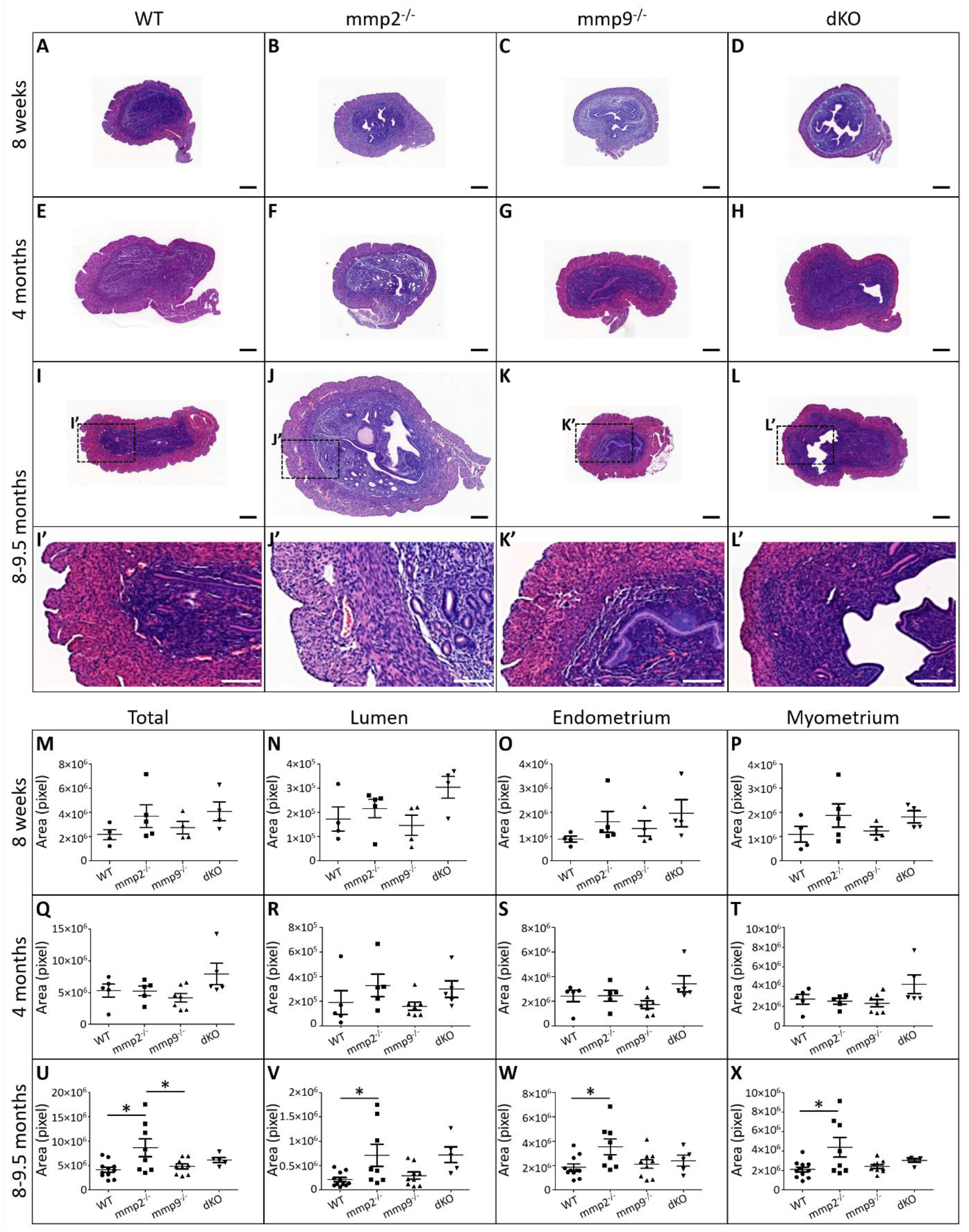
Uterine histology reveals enlargement of the overall uterine tissue in mmp2^-/-^ at 8-9.5M of age. (A-H) Histological sections from uterine horn demonstrate small differences in single- or double-KO samples compared to WT, at the age of 8w and 4M. (I-L) Significant increase in overall tissue size was found at 8M old mmp2^-/-^ uteri compared to mmp9^-/-^, dKO and WT. (I’-L’) higher magnification of the marked area in I-L. (M-T) total area, and lumen, endometrium and myometrium area were found to be similar between the different genotypes at the age of 8w and 4M. (U-X) Specific measurements and statistical analysis show that at 8-9.5M, mmp2^-/-^ uteri demonstrate a significant enlargement of all measured elements. n=4-12 for each genotype and age. At A-L’, bar=200µm.

Notably, in concomitant with the relatively low proportion of the dystocia occurrence in mmp9^-/-^ females (9.23%, Fig. 1), the histology of these uteri did not present major differences compared to WTs’ in all examined ages, in terms of the total area of the tissue, or the specific area of the myometrium, endometrium and lumen. Furthermore, the histological characteristics of the dKO uteri, which lacked both MMP2 and MMP9, did not demonstrate a severer phenotype compared to that of mmp2^-/-^ at any ages (Fig. 3, compare D, H and L to B, F and J and to A, E and I, respectively). Additionally, the specific measurements of the dKO histology demonstrated that while the total area and the myometrium, endometrium and lumen areas of mmp2^-/-^ differed from that of WT significantly (Fig. 3 U-X), the same parameters measured for dKO, did not differ from neither mmp2^-/-^ nor WT. This result, which is in agreement with the less-prominent percentage of dystocia observed in dKO females compared to mmp2^-/-^ females (Fig. 1), strongly suggests that the additional loss of *mmp9* on the background of *mmp2* loss (in dKO and mmp2^-/-^, respectively), activates some compensative mechanisms that rescue the phenotype, at least to some level.

### Mmp2^-/-^ nulliparous uterine at 12M demonstrates signs of myometrial fibrosis

As MMP2 has been previously shown to degrade collagen as one of its substrates [33,34], and as collagen accumulation is associated with fibrosis in different tissues [35,36], we next asked whether the observed pathophysiology in the MMP2-nulls is coupled with accumulation of collagen in the uterine wall. Masson’s Trichrome staining for collagen fibrils was performed on transverse sections from nulliparous WT and mmp2^-/-^ uteri at 12M of age, demonstrating an increase in collagen fibrils in mmp2^-/-^ tissue as shown by the increased blue staining mostly in the circular myometrium layer, compared to WT (Fig. 4, compare B-B’ to A-A’; arrows). This result suggests that one of the underlying mechanisms of the disrupted functionality of the uterine tissue during parturition in the *mmp2* KO, is due to insufficient remodeling of the uterine myometrium which results in fibrotic myometrium. This fibrosis can be the cause for insufficient contractions of the uterus during parturition, which in turn lead to the defective parturition process and dystocia.

**Figure 4.**
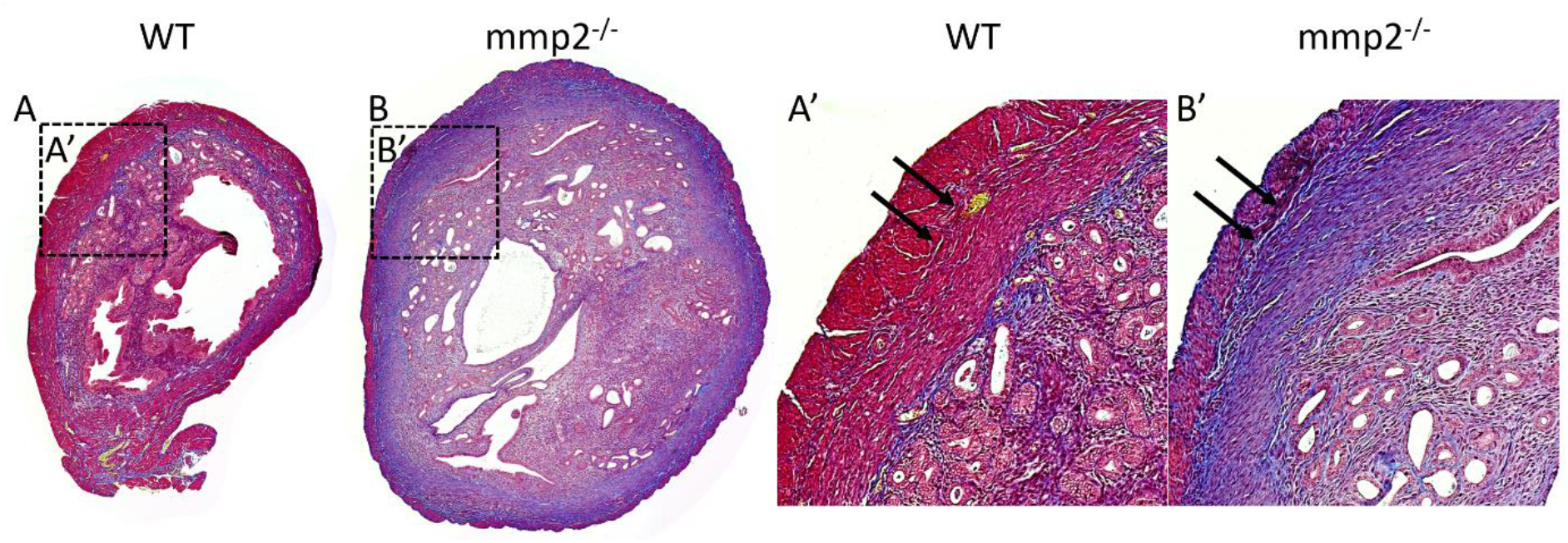
Masson’s trichrome staining demonstrate extensive fibrosis in 12M old mmp2^-/-^ nulliparous uterine tissue. (A-B) Transverse sections from 12M old WT or mmp2^-/-^ nulliparous uteri stained with Masson’s Trichrome to recognize collagen fibrils, respectively. (A’-B’) higher magnification of marked area in A and B, respectively, demonstrating increase in collagen fibrils in mmp2^-/-^ uterus (arrows).

## Discussion

In this study we investigated the role of the gelatinases MMP2 and MMP9 in uterine function during parturition process using a loss-of-function approach to genetically knock out MMP2, MMP9 or both in the mice model. We found that while *mmp9*-loss is only affecting partially the parturition process, *mmp2*-loss leads to a severe defective parturition which included very high rates of dystocia when reaching to term. We demonstrated that 31.37% of all mmp2^-/-^ pregnancies result in dystocia, in which females present one or more (or none) living newborns together with one or more dead fetuses still in the uterus. This percentage is very low in WT females (1.01%) and much less high in mmp9^-/-^ females (9.23%). Surprisingly, the dystocia proportion increases in mmp2^-/-^mmp9^+/-^ females to 40%, but decreases to 21.95% in the dKO. Furthermore, when the histology of the nulliparous uterine was examined at several ages, it showed a significantly enlarged tissue in mmp2^-/-^, rather than in mmp9^-/-^ or dKO as compared to WT, with increased tissue area of the myometrium, endometrium, and lumen.

Importantly, Masson’s Trichrome staining for collagen fibrils, demonstrated deposition of more collagen in the mmp2^-/-^ uterine tissue compared to WT at the age of 12M. As MMP2 is largely known for its proteolytic activity to degrade collagen (among other ECM molecules) [34,37,38], it is reasonable to speculate that when this activity is lost, it results in accumulation of collagen in the uterine ECM, which in turn causes myometrial fibrosis. Our study provides data that support this conclusion, suggesting that the increased fibrosis in nulliparous mmp2^-/-^ uterine may be one of the causes for insufficient contractions during parturition, ending with dystocia and in-uterus fetal death. Furthermore, while during pregnancy the cervix must be firm so that the fetus is not prematurely expelled, by reaching to term, the cervical softening is initiated by cascade of events that includes remodeling in the ECM, making the cervix more compliant to the parturition process [1]. Based on the suggested fibrosis in the uterine myometrium of mmp2^-/-^ females, it is possible that also the cervix of these females fails to properly remodel the ECM which leads to cervical fibrosis. In agreement with our results, previous studies have found that the KO mice for Anthrax toxin receptor 2 (*Antxr2*) present defective parturition and dystocia, which was suggested to be the result of extensive fibrosis in the uterus and cervix [39,40]. The researchers demonstrated an aberrant deposition of ECM proteins such as type I collagen, type VI collagen and fibronectin, together with a marked disruption of both the circular and longitudinal myometrial cell layers. Notably, in the uterine of Antxr2^-/-^ females, a decrease in the active form of MMP2 was demonstrated and the researchers provided supporting information for a possible mechanism by which ANTXR2 regulates the activity of MMP14 which further activates MMP2. This study reinforces our data supporting an important role for MMP2 in uterus ECM remodeling. Furthermore, a different study found in Geranylgeranyl pyrophosphate synthase (*Ggps1)* conditional KO (cKO), that 75% of all pregnancies ended with dystocia [41]. The researchers isolated uterine muscle strips from adult nonpregnant mice and found that the spontaneous contraction rate and amplitude in *Ggps1* cKO mice was largely decreased compared to controls. Since the Rho/Rho-associated protein kinase (Rock) pathway is associated with smooth muscle contraction, they found the membrane association of RhoA significantly decreased after *Ggps1* deletion in the myometrium, and the expression of Rock2, the downstream target of Rho, was also decreased. Thus, they concluded that *Ggps1* deletion disrupts the RhoA/Rock2 pathway, causing uterine contraction and parturition problems. Interestingly, Rho has been found to regulate MMP expression at various cell types [42–44], raising the possibility that they are also co-involved in governing parturition process.

Cephalopelvic disproportion (CPD) is one of the leading causes for dystocia in women [45,46]. CPD occurs when there is mismatch between the size of the fetal head and size of the maternal pelvis, resulting in “failure to progress” in labor for mechanical reasons; if untreated, the consequence is obstructed labor that can endanger the lives of both mother and fetus [45,46]. Hence, an additional possible mechanism for the abnormal parturition and dystocia can be related to the defective skeleton of mmp2^-/-^ mice [21]. We have recently uncovered the role of MMP2 and MMP9 in skeleton development and found that, while MMP9 participates in endochondral ossification process and its loss affects the length of long bones, MMP2 participates in intramembranous ossification process and its loss results in shorter but wider skulls of the neonates together with thinner and less mineralized cortices [21]. Since also the pelvic bones are composed by irregular and flat bones which develop through intramembranous ossification [47–49], we cannot rule out that, that similar to the skull, the pelvic bones of mmp2^-/-^ mice are mal-developed resulting in an abnormal pelvis. Thus, impaired pelvis of the mmp2^-/-^ pregnant female along with the wider skull observed in the mmp2^-/-^ fetuses, can be one of the possible causes for cephalopelvic disproportion resulting in dystocia in these mice, as was previously reported in women [45,46,50]. Moreover, as we found that while some P0 mmp2^-/-^ newborns have significantly wider skull, others have milder or no skull defect [21], this can explain why some fetuses are able to be expelled from the uterus while others get physically stacked and die.

Finally, the use of knockout mice for different MMPs throughout the years has allowed the investigation into their role in various aspects of physiology, including reproductive function. For example, Sternlicht and Werb have summarized the non-malignant phenotypes associated with the genetic modifications of MMP1, MMP2, MMP3, MMP7, MMP9, MMP11, MMP12, and MMP14, along with TIMP1, TIMP2, and TIMP3 [51]. However, none of these reports showed major impact on the reproductive axis in the initial characterization of these genetically modified animals. Specifically, deletion of MMP9 or TIMP1 was not associated with an obvious reproductive phenotype in the initial description of these mice [27,52], but subsequent analysis has revealed changes in the reproductive axis [23,53]. For example, TIMP1-null females achieved pregnancy at a lower rate than wildtype mice (52% *vs* 78%) and had significantly fewer pups per litter [23]. As TIMP activity was found to be closely linked to MMP2 activation in fibrosarcoma cells *in vitro*, further studies are needed to explore whether MMP2 activity is impaired in the uterine of the TIMP1-nulls.

A confounding factor in trying to understand MMP’s action/role in a given physiological or pathological condition, is that removal of one MMP often results in compensation by other members of the MMP family. For example, the deletion of MMP7 results in a 10–12-fold increase in MMP3 and MMP10 during uterine involution [54]. Furthermore, we showed in embryonic NCCs that the loss of one gelatinase (MMP2 or MMP9), is compensated by the other one, leading to normal migration phenotype of the NCCs [18]. Furthermore, we have recently showed by RNAseq analysis that the loss of both gelatinases in the bone tissue leads to an increase in other MMPs’ transcript, such as *mmp13, mmp14* and *mmp15* [21]. Hence, it is possible that also in reproduction mechanism the genetic loss of one MMP is compensated by other MMPs, preventing the occurrence of a specific fertility-related phenotype in experimental animal models. Thus, the different results observed in mmp2^-/-^ and mmp9^-/-^ females can be explained by a unilateral compensation mechanism between the two gelatinases, by which *mmp2* can rescue to some level the loss of *mmp9*, but not *vice versa*; hence, the remaining of *mmp9* allele in mmp2^-/-^ cannot rescue the defective parturition phenotype. This suggested mechanism can also explain the higher rate of dystocia when only one additional allele of *mmp9* is missing, in mmp2^-/-^mmp9^+/-^, to 40%. Nevertheless, the result of the dKO, presenting lower rates of dystocia (21.95%), suggests that the mechanism is more complexed than that and needs to be further investigated.

In summary, our data demonstrated that mmp2^-/-^ female mice suffer from defective parturition process which ends in more than 30% of cases with dystocia and the inevitable death of both fetuses and pregnant females. Our data highlights that this percentage is increased to 40% in the lack of one allele of the other gelatinase, MMP9 (mmp2^-/-^mmp9^+/-^) but decreases to 21.95% in dKO. Our histological analysis hints for a mechanism by which mmp2-loss leads to increases fibrosis in the mmp2^-/-^ myometrium tissue, compared to WT. These findings calls for further analysis on the roles and mechanism of action of gelatinases in the prevention of dystocia.

